# Methyl donor deficient diets cause distinct alterations in lipid metabolism but are poorly representative of human NAFLD

**DOI:** 10.1101/162131

**Authors:** Marcus J Lyall, Jessy Cartier, James A Richards, Diego Cobice, John P Thomson, Richard R Meehan, Stephen M Anderton, Amanda J Drake

## Abstract

Non-alcoholic fatty liver disease (NAFLD) is a global health issue. Dietary methyl donor restriction is used to induce a NAFLD/non-alcoholic steatohepatitis (NASH) phenotype in rodents, however the extent to which this model reflects human NAFLD remains incompletely understood. To address this, we undertook hepatic transcriptional profiling of methyl donor restricted rodents and compared these to published human NAFLD datasets.

Adult C57BL/6J mice were maintained on control, choline deficient (CDD) or methionine/choline deficient (MCDD) diets for four weeks; the effects on methyl donor and lipid biology were investigated by bioinformatic analysis of hepatic gene expression profiles followed by a cross-species comparison with human expression data of all stages of NAFLD.

Compared to controls, expression of the very low density lipoprotein (VLDL) packaging carboxylesterases (*Ces1d*, *Ces1f*, *Ces3b*) and the NAFLD risk allele *Pnpla3* were suppressed in MCDD; with *Pnpla3* and the liver predominant *Ces* isoform, *Ces3b*, also suppressed in CDD. With respect to 1-carbon metabolism, down-regulation of *Chka*, *Chkb*, *Pcty1a*, *Gnmt* and *Ahcy* with concurrent upregulation of *Mat2a* suggests a drive to maintain S-adenosylmethionine levels. There was minimal similarity between global gene expression patterns in either dietary intervention and any stage of human NAFLD, however some common transcriptomic changes in inflammatory, fibrotic and proliferative mediators were identified in MCDD, NASH and HCC.

In conclusion, this study suggests suppression of VLDL assembly machinery may contribute to hepatic lipid accumulation in these models, but that CDD and MCDD rodent diets are minimally representative of human NAFLD at the transcriptional level.

**Summary statement:** We used transcriptional profiling of methyl donor restricted rodents to examine effects on methyl donor and lipid biology. We report novel mechanisms for lipid accumulation in this model and describe significant disparity between both dietary interventions and human disease.

## Introduction

Non-alcoholic fatty liver disease (NAFLD) is the predominant cause of chronic liver disease in the developed world with an estimated prevalence of between 20-68% ^1^. The accumulation of hepatic fat in the form of triglycerides and other lipid species in NAFLD has two major clinical consequences. Firstly, a subgroup of patients with hepatic steatosis will progress to an inflammatory hepatitis, hepatic cirrhosis and in some cases hepatocellular carcinoma (HCC) ^2^. Secondly, almost all patients with NAFLD also exhibit hepatic insulin resistance, which can associate with impaired glucose uptake, increased gluconeogenesis and type 2 diabetes, possibly as a direct consequence of the increased hepatic lipid load ^1,3,4^. Together these conditions are responsible for significant morbidity and mortality and represent a substantial burden for health resources ^5^.

The molecular mechanisms underpinning NAFLD pathology are incompletely understood and as such there is a need for accurately representative rodent models in which to investigate this common disease and to trial novel therapeutics. Given the association of NAFLD with human obesity, the use of high fat diet feeding in rodents remains a popular model in which to investigate mechanisms. However whilst high fat feeding generates a NAFLD-like picture, the disadvantages with this model include the protracted time required to induce even mild non-alcoholic steatohepatitis (NASH), the lack of malignant transformation to HCC even with prolonged exposure, and the variation in the histological and transcriptional changes due to the behavioural characteristics of mice in social groups^6–8^. Thus, a number of alternative models have been employed. In rodents, dietary restriction of the methyl donors methionine and/or choline rapidly and reliably induces a spectrum of liver injury histologically similar to human NAFLD, within weeks of instigation ^9,10^. Although the precise biological mechanisms responsible for the predictable phenotypic changes are poorly understood, the histological similarity to human steatosis (choline deficient diets; CDD) and NASH (methionine and choline deficient diets; MCDD) means that these models have been used in mechanistic and therapeutic studies for a number of years ^11–13^. Since impaired metabolism of the key methyl donor S-adenosylmethionine (SAMe) is a well documented feature of chronic liver disease regardless of aetiology ^14–16^, there may be common molecular mechanisms which may present an opportunity for the rapeutic intervention. Although a number of transcriptional changes have been reported during the progression of human NAFLD, no detailed transcriptional comparisons have been performed to identify similarities or differences between human disease and the CDD and MCDD models of NAFLD, despite their widespread use. In this study we set out firstly to dissect potential mechanisms underpinning the development of liver pathology in CDD and MCDD models by mapping pathways of lipid and one-carbon metabolism, and secondly to evaluate their potential usefulness as models of human disease. To address these aims we have examined in detail the transcriptional profiles in liver from mice maintained on CDD and MCDD and compared these with published human NAFLD transcriptome data series.

## Materials and Methods

### Animals

All experiments were carried out under a UK Home Office licence and with local ethical committee approval. Adult C57BL/6J mice (Charles River, Tranent, UK) were maintained under controlled conditions in social groups of 5 animals per cage. A 12-hour light cycle (07.00h to 19.00h) and twelve hour dark cycle was implemented throughout. The temperature was maintained at 22°C +/− 2°C. Mice were maintained on control, CDD or MCDD diets (Dyets, Bethlem, PA) for 4 weeks and then killed and tissues collected and used for histology or snap-frozen and stored at −80C. Diet composition can be found in Supplementary Table 1.

### Histology staining, triglyceride and SAMe quantification

Livers were removed and sections were fixed in methacarn solution (methanol:chloroform:glacial acetic acid; ratio 6:3:1) and mounted in paraffin blocks prior to staining with haematoxylin and eosin or picosirius red. Hepatic triglyceride concentration was determined by spectrophotometric analysis (BioVision, Milpitas, USA) as previously described ^17^. Image analysis for fibrosis content was performed in ImageJ (http://imagej.nih.gov/ij/). SAMe was quantified by matrix-assisted laser desorption ionization mass spectrometry imaging (MALDI-MSI) using the 12T SolariX MALDI-FTICR-MS (Bruker Daltonics, MA, US) as previously described ^18^.

### Reverse transcription and qPCR

RNA was extracted from snap frozen liver tissue using the RNeasy kit (Qiagen, Manchester, UK). 800ng of RNA was DNAse treated using Promega RQ1 DNAase (Promega, Southampton, UK) and reverse transcribed with the High Capacity cDNA Reverse Transcription Kit (Life Technologies, Paisley, UK). Quantitive real time PCR was performed using Roche Universal Probe Library assays or TaqMan qPCR assays (Life Technologies, Paisley, UK) (Supplementary Table 2) using the Roche Lightcycler 480 and associated software (Roche, West Susssex, UK). Gene expression is displayed relative to mean of three housekeeping genes (Gapdh, Ppia, Ldha).

### Transcript analysis

RNA labelling was performed on 500ng RNA using the Illumina Total Prep RNA amplification kit (Life Technologies. Paisley, UK) and subsequently hybridised to Illumina Mouse-ref6 expression bead arrays as per the manufacturer’s instructions, at the Edinburgh Clinical Research Facility, Western General Hospital, Edinburgh, UK. Intensity data were generated using a HiScan array scanner (Illumina, San Diego, USA) and analysed using iScan Illumina software. Data analysis and generation of plots were perfomed in RStudio (http://www.rstudio.com) with R version 3.1.2. Data import, quality control, normalisation and between array adjustment was performed using Lumi package and differential expression was determined using Limma package (Bioconductor.org). Unsupervised clustering was performed using Euclidean distance. Where multiple probes mapped to the same gene, the median result was used. Data have been uploaded to EBI-Array Express, accession number E-MTAB-3943.

### Pathway Analysis

Gene Ontology and pathway enrichment was performed using the GOstats package (Bioconductor.org). Investigation of fat handling was performed by interrogating relevant pathways of lipid metabolism and insulin signalling from the Kyoto encyclopedia for genes and genomes (KEGG) module database (http://www.genome.jp/kegg/module.html). KEGG module sets ‘M00003 gluconeogenesis’, ‘M00086 beta-Oxidation, acyl-CoA synthesis’, ‘M00083 Fatty acid biosynthesis, elongation’ and ‘mmu_M00089 Triacylglycerol biosynthesis’, ‘mmu00071 Fatty acid degradation’, and KEGG pathways ‘Fat Digestion and Absorption’ and ‘Insulin Signalling Pathway’ and ‘Glycerolipid Metabolism’ were used. In addition, the family of carboxylesterase (Ces) genes were analysed due to their recently discovered role in triglyceride hydrolysis ^19–21^. Finally, to dissect the link between lipid metabolism and one carbon metabolism, relevant mediators were analysed and mapped to known biochemical pathways.

### Cross species comparison

A comparison of transcriptional data from both CDD and MCDD was made with published human expression sets of normal liver, simple steatosis and NASH (GSE48452, E-MEXP-3291), and HCC (GSE638980) ^22–24^. Data sets were retrieved from the ArrayExpress archive (http://www.ebi.ac.uk/arrayexpress/). Predicted gene orthologues were determined using *homologene* via the Hugo Gene Nomenclature Committee server (http://www.genenames.org). For all human NASH, HCC and all mouse data sets, a transcriptional threshold of 1.5 with an adjusted p value of <0.05 was applied. For human steatosis gene sets alone, the fold change transcriptional threshold was reduced to 1.2 to allow comparison with the relatively mild transcriptional derangement observed. All genes found to be dysregulated at each stage of human NAFLD were then examined in the CDD and MCDD data sets to determine if expression was altered, and if so,, what was the direction of change. Genes dysregulated in human and mouse models were then depicted in scatter plot analyses with linear regression used to compare data sets.

### Statistics

Animal model and qPCR statistical analysis was performed using Prism GraphPad software (GraphPad Software Inc.). Data were routinely analysed for outliers, normalisation and sphericity where required. Non-parametric data were either log transformed or a non-parametric test used as indicated.

## Results

### Phenotype

Mice on CDD gained significantly less weight than animals on a control diet whereas MCDD fed mice lost weight from the outset, consistent with previous observations ^17,25^ (Figure 1A). At the end of the experiment, hepatic triglyceride content was significantly higher in both CDD and MCDD groups compared to controls, but there was no significant difference between the two interventions (Figure 1B). MCDD liver weights were lower in MCDD fed mice but not CDD (Figure 1C). Histological analysis revealed severe hepatic steatosis in both groups (Figure 1D) with a significant increase in hepatic fibrosis in the MCDD group (Figure 1E).

**Fig. 1.**
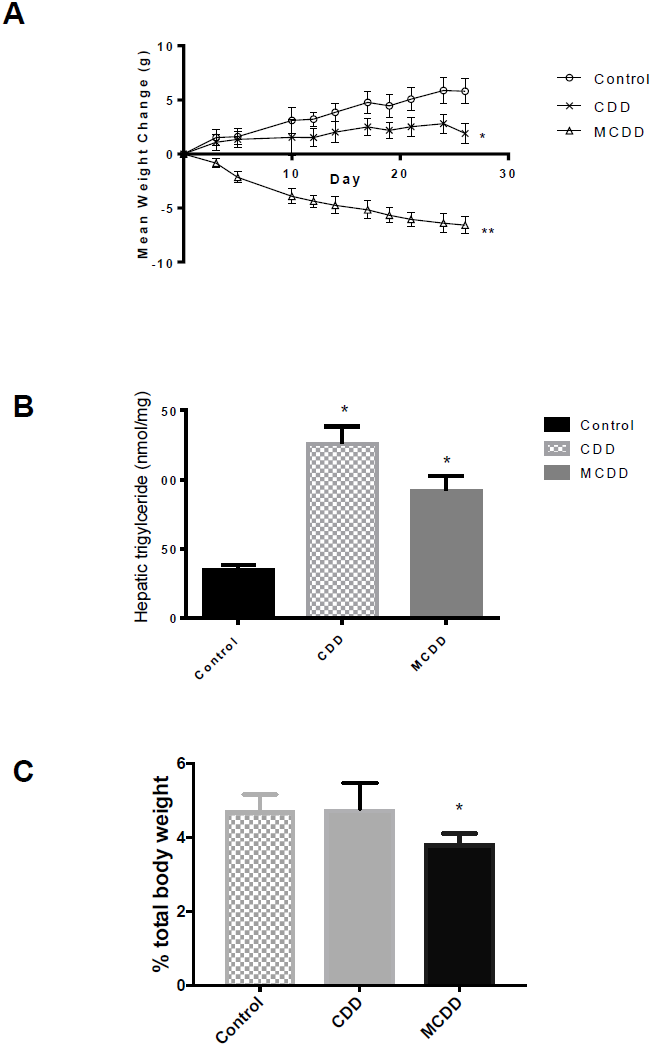
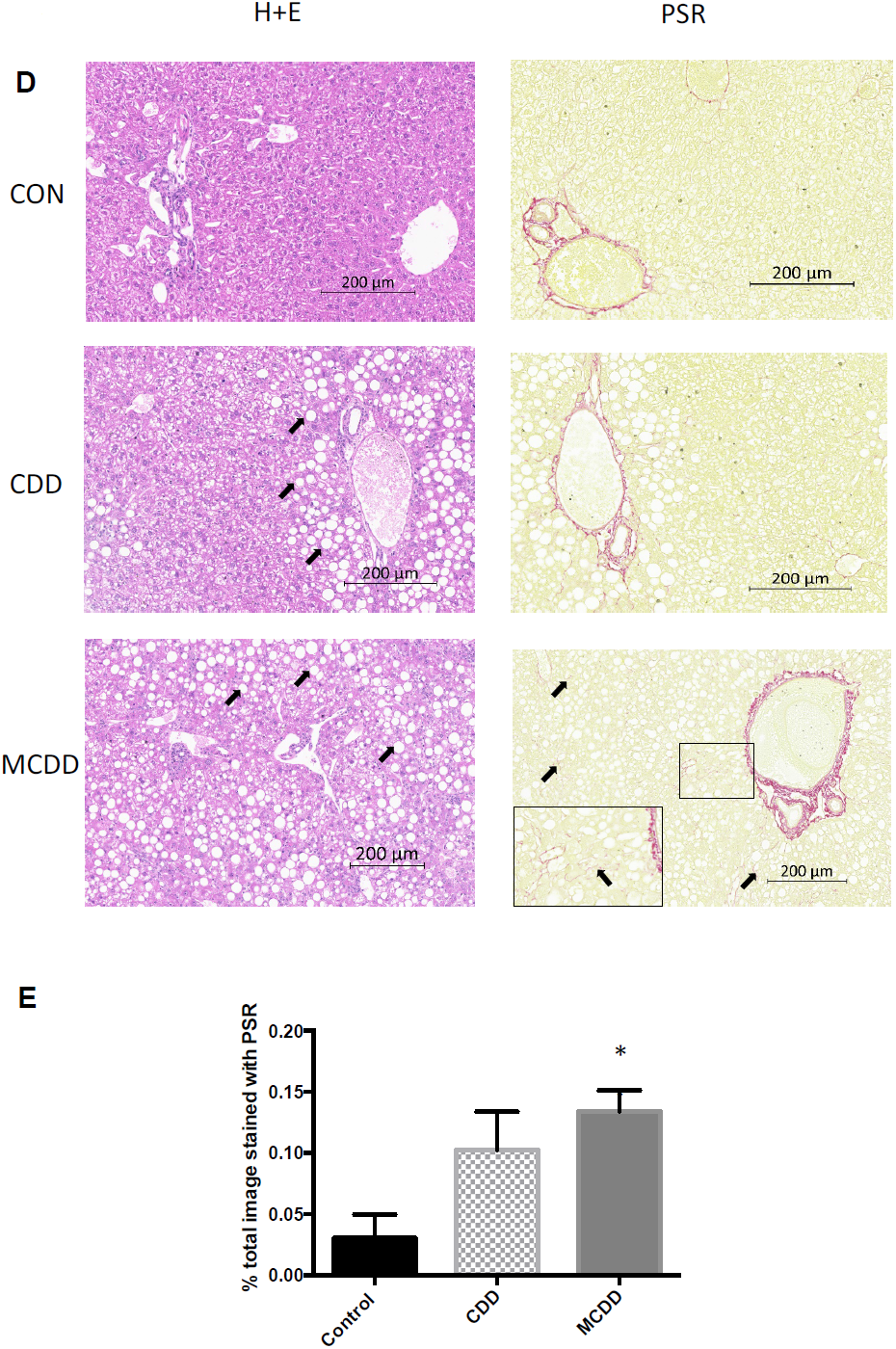
Body weight and hepatic lipid content. A) In comparison with control animals, mice on CDD gain less weight whereas those on MCDD lose weight. B) Hepatic liver triglyceride content was increased on both CDD and MCDD diets. C) Liver weight was reduced in MCDD fed mice but not CDD. D) Liver histology from control, CDD and MCDD mice. H+E = Hematoxylin and Eosin staining showing marked macrovesicular steatosis in both CDD and MCDD mice (white arrows). PSR = Picosirius Red stain staining for new collagen formation showing increased periportal and interstitial fibrosis in MCDD animals (black arrows). In MCDD liver stained with PSR, insert at higher magnification shows increased fibrosis more clearly. E) Quantified fibrosis content (PSR positive staining as a percentage of entire image). n = 10 per group for all figures. * P < 0.05 ** P < 0.01 (one way ANOVA with Tukey post hoc test versus control animals). Error bars = +/− SEM

### Gene Expression Changes

Next we carried out analysis of the transcriptome in control, CCD or MCDD mouse livers, each n=4. This approach allowed us to interrogate ~18,000 transcripts per mouse liver and analysis of total datasets revealed a number of transcriptional differences between animals. Unsupervised clustering of the 500 most variable transcripts between all animals was sufficient to cluster into the different dietary interventions (Figure 2A). Both interventions induced a >1.5-fold differential expression in multiple transcripts when corrected for multiple testing (adjusted P value <0.05, Benjamini-Hochberg test) (Figure 2B-D). The top 100 up- and down-regulated genes in each group are shown in Supplementary Tables 3 and 4. Of the 234 genes differentially expressed in CDD, 194 (82.4%) were also differentially expressed in MCDD. The additional restriction of methioine over and above choline induced the differential expression of a further 1032 transcripts (Figure 2B).

**Fig. 2.**
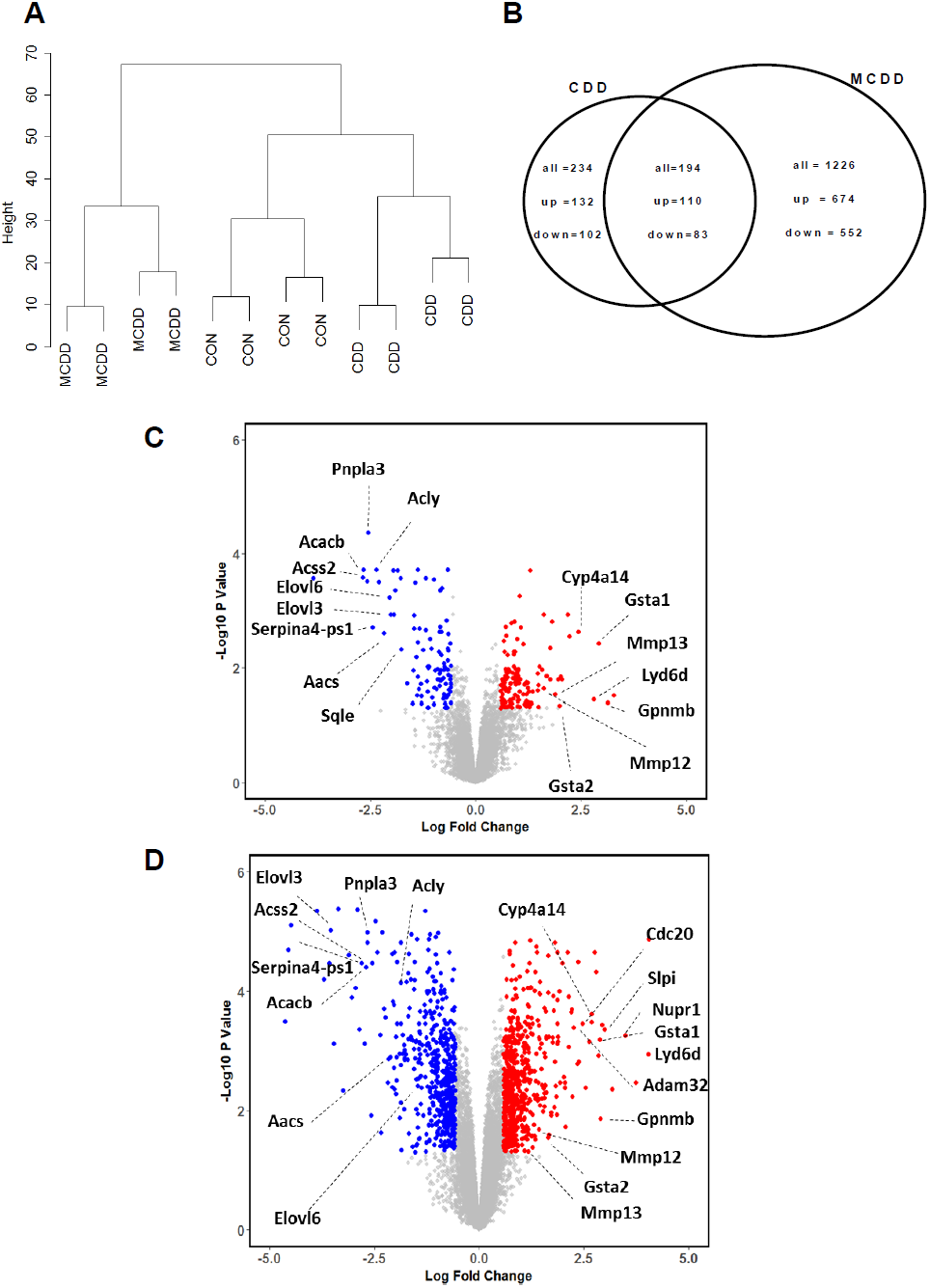
Differential gene expression. A) Transcript profiles of each dietary intervention were sufficiently consistent to cluster by Euclidean distance. B) Venn diagram demonstrating high degree of overlap in dysregulated transcripts (≥2 fold change) in each group and the direction of transcriptional change. C and D) Volcano plots demonstrating transcriptional changes in CDD (C) and MCDD (D) mice. Dotted lines and blue colour represent adjusted P values < 0.05 and two fold differential expression change.

Transcripts showing at least a 2-fold change were segregated into up-regulated and down-regulated gene lists and examined for over-representation within all detected transcripts on the array platform. GO-terms for lipid, sterol, fatty acid and organic acid biosynthesis were markedly over-represented in the list of suppressed genes. Over-expressed pathways in CDD mice included ‘immune system process’ and ‘inflammatory response’. Up-regulated pathways in mice on the MCDD diet included ‘positive regulation of mitotic cell cycle’ and ‘negative regulation of cell cycle arrest’ (Supplementary Figure 1).

The most up-regulated genes in CDD mice included immune mediators (*Gpnmb*, *Ly6d*); fibrosis mediators (*Mmp12*, *Mmp13*); and the detoxification enzymes *Gsta1* and *Gsta2* and the microsomal enzyme *Cyp4a14*. The most suppressed genes in CDD included lipid synthesis genes (*Sqle*, *Elovl3*, *Elovl6*, *Aacs*, *Acly*, *Acss2*, *Acacb*), the endopeptidase inhibitor *Serpina4-ps1* and the multifunctional triglyceride metabolism enzyme *Pnpla3*. Whilst these genes were similarly differentially expressed in MCDD mice, volcano plots revealed considerably more severe transcriptional derangement in MCDD compared with CDD, both in the number and fold change of differentially expressed genes (Figures 2C and 2D). Additional genes up-regulated in MCDD included the mitotic proteins *Cdc20*, *Nupr1*, the metalloproteinase *Adam32*, the inflammatory mediator *Slpi* and the aldo-ketoreductase *Akrb7*.

qPCR validation of array findings was performed for known mediators of lipid uptake (*Lpl*), putative contributary genes to hepatic fat accumulation in NAFLD (*Scd1*, *Aacs*, *Fasn*, *Mlxipl*, *Acsl1*) and hepatic fibrogenesis (*Mmp12*), and *Pdk1* which is an important link between insulin signaling and HCC. In addition we analysed gene expression of four members of the *Ces* family due to their functional role in triglyceride hydrolysis. Gene expression by qPCR was consistent with array findings for all genes (Figures 3A and B).

**Fig. 3.**
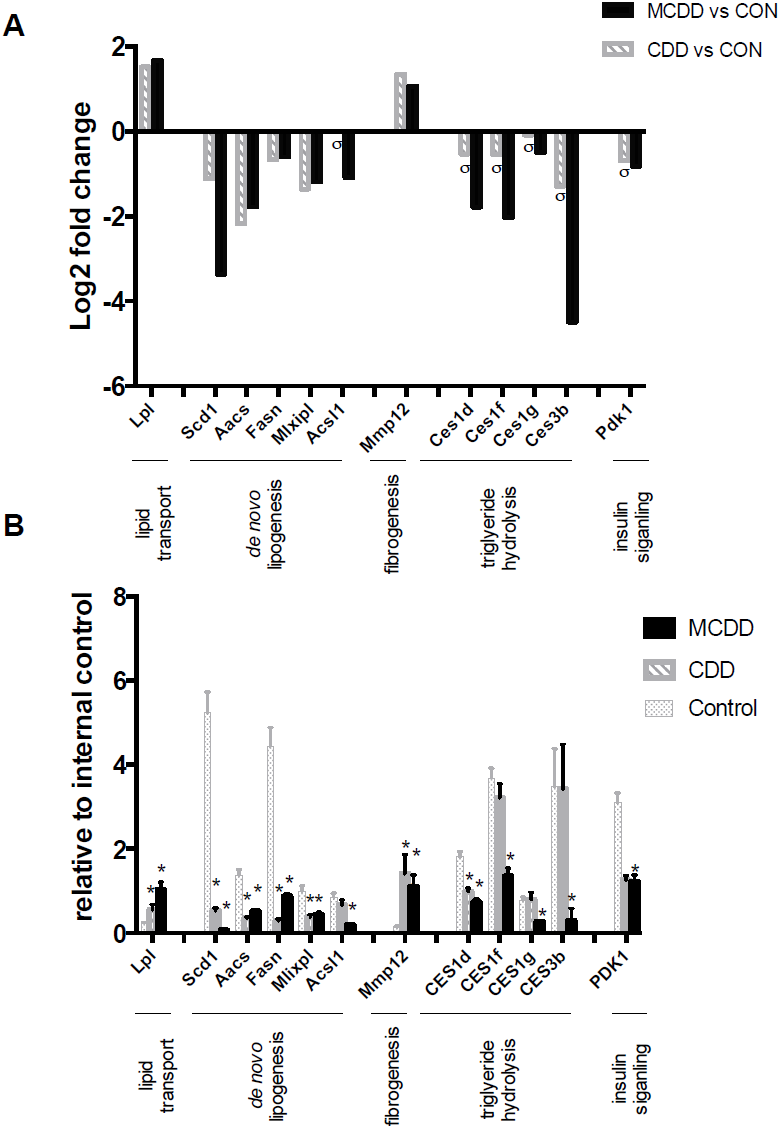
CDD and MCDD induce changes in the hepatic expression of genes important in lipid metabolism and storage. (A) Microarray analysis and (B) qPCR validation of selected genes important in lipid transport, *de novo* lipogenesis, fibrogenesis, triglyceride hydrolysis and insulin signalling. (A) Adjusted P value < 0.05 (FDR) for all samples except those marked σ. (B) * = P < 0.05 versus control, one-way ANOVA with Bonferroni post hoc analysis. Error bars = +/− SEM

### Lipid Pathway Analysis

We then proceeded to examine pathways of lipid uptake, synthesis and disposal. Differentially expressed genes in relevant KEGG pathways in either group are depicted in Figure 4. Expression changes were generally greater in MCDD than CDD although they occurred in the same direction. Previous studies have suggested an increase in lipid uptake with MCDD, with upregulation of some of the FATP/solute carrier family 27 genes in association with increased sequestration of isotope labelled fatty acids ^17,26^, however we noted only the down-regulation of the fatty acid translocase Scla27a5 (FATP5) with no change in the other FATP isoforms. We did identify marked upregulation of lipoprotein lipase, a key mediator in triglyceride hydrolysis from lipoproteins. In keeping with the gene ontology analysis, the global picture suggests suppressed hepatic lipid synthesis. Perturbed genes in pathways of fatty acid biosynthesis initiation, Acyl-CoA synthesis, fatty acid elongation and cholesterol synthesis were almost universally down-regulated.

**Fig. 4.**
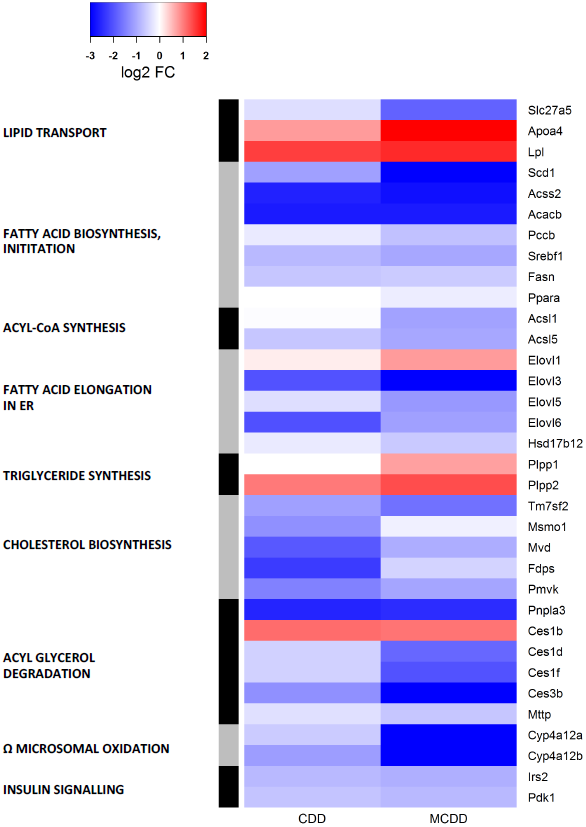
Heatmap depiction of expression changes in KEGG pathways of lipid transport, lipid synthesis and degradation and insulin signalling. Up- and down-regulated genes in each dietary intervention versus control animals are demonstrated by colour key. Pathways of lipid and cholesterol synthesis are globally suppressed.

We then examined dominant pathways of hepatic fatty acid fate (triglyceride synthesis, β-oxidation, oxidation and peroxisomal oxidation). These were found to be relatively unaffected, apart from an upregulation of the phospatidic acid phosphatases *Pap2a* and *Pap2c* (which convert phospatidic acids to diacylglycerol) and the peroxisomal fatty acid elongation enzyme Elovl1 in MCDD. The Cyp4a family of enzymes broadly catalyse the microsomal (ω) oxidation of saturated and unsaturated fatty acids and have reported to be upregulated by dietary and drug-induced hepatic inflammation ^27–29^. Interestingly, isoforms of Cyp4a enzymes demonstrated marked bi-directional differential expression with Cyp4a12a and Cyp4a12b strongly suppressed in CDD and MCDD whilst Cyp4a14 was upregulated by 4-fold with both diets. Other microsomal oxidation Cyp450 isoforms (Cyp4a10, Cyp4a32, Cyp4a29, Cyp4a30b) were unchanged. Cyp2e, which has previously been reported to be upregulated in MCDD was also unchanged ^29^. In keeping with the theory of impaired very-low-density lipoprotein (VLDL) secretion in NAFLD ^17,30^, endoplasmic reticulum (ER)-associated mediators of triglyceride hydrolysis and VLDL assembly (*Pnpla3, Mttp* and the carboxylesterase enzymes: *Ces1d*, *Ces1f*, *Ces3b*) were markedly suppressed in MCDD, with Pnpla3 also suppressed in CDD. Interestingly, expression of the *Ces1b* isoform was clearly up-regulated in both groups, in contrast to the other members of this class. Apoa4, a lipid binding protein involved in the expansion and secretion of VLDL particles was significantly induced. Finally, two key intermediaries in hepatic insulin signal transduction (*Irs2* and *Pdk1*) were also down-regulated.

### One Carbon Metabolism

Given the importance of choline and methionine as methyl donors, we then proceeded to examine the expression of genes important in one-carbon metabolism. There were striking changes in the expression of genes associated with choline, methionine and phosphatidylcholine (PC) metabolism in both interventions, broadly in the same direction (Figure 5A). Two pathways demonstrated significant down-regulation of key enzymes in MCDD: including genes important in the synthesis of PC from choline (*Chkb*, *Pcyt1a*) and in the conversion of SAMe to homocysteine (*Gnmt*, *Ahcy*). Furthermore, the expression of enzymes that contribute to the clearance of methionine, SAMe and S-adenosylhomocysteine (*Mthfd1*, *Gnmt*, *Achy*, *Dnmt3b*) were suppressed in both groups. In MCDD mice, there was also marked upregulation of expression of *Mat2a*, an enzyme necessary for the synthesis of SAMe from methionine. In the light of these results, we measured SAMe concentrations in each group using MALDI-MSI. This demonstrated a significant reduction in SAMe in MCDD mice with no change in CDD (Figure 5B). A summary of pathways showing changes in enzyme expression is shown in Figure 5C.

**Fig. 5.**
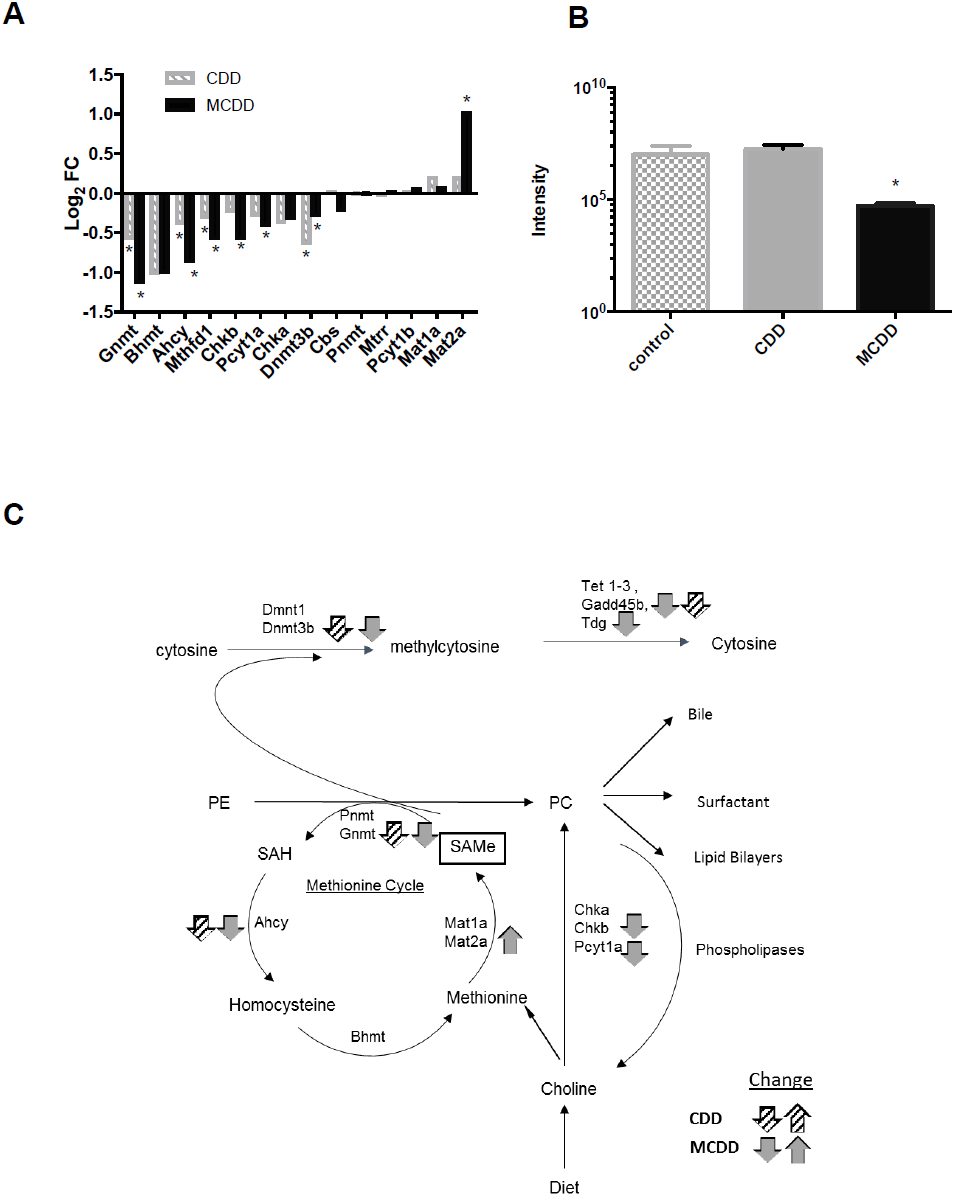
CDD and MCDD induce changes in the hepatic expression of genes important in one-carbon metabolism. (A) Transcriptional changes in one-carbon metabolism enzymes induced by CDD and MCDD diets (* = adjusted P value < 0.05, Benjamini – Hochberg analysis). (B) SAMe levels as measured by MALDI analysis. (* = P< 0.01 one-way ANOVA with Bonferroni post hoc analysis). C) Metabolic interactions between methionine cycle, phosphatidylcholine synthesis and DNA methylation. Grey (CDD) and black (MCDD) arrows depict expression changes in each pathway. PE=phosphatidylethanolamine, PC=phosphatidylcholine, SAH=S-adenosylhomocysteine, SAMe = S-adenosylmethionine. Adapted from Li and Vance 2008.

### Comparison With Human NAFLD Data Sets

We then proceeded to compare our transcriptional findings with large published expression sets of three stages of NAFLD: simple steatosis, NASH and HCC ^22–24^ (Figure 6). While a number of microarray studies have been performed in NAFLD ^31–33^ we selected data sets from Ahrens *et al* and Lake *et al* (GSE48452, E-MEXP-3291) due to the detailed patient and histological descriptors (including Kleiner NAFLD activity (NAS) score) confirming NAFLD stage ^22,23^. In addition, these data sets include all three NAFLD stages and control samples in the same data series reducing assay variation and are directly available from the ArrayExpress repository. This allowed direct comparison of obese subjects with simple steatosis (n=22, NAS score <3), and patients with NASH (n = 24, NAS score 3-5) with well characterised controls (n = 37). Details of subject numbers and arrays used are in Supplementary Table 5. There are no current data sets available from exclusively NAFLD-induced HCC. We therefore used a large data set of mixed Hepatitis C and alcohol-induced HCC samples (n=228), which are directly compared with cirrhotic liver samples (n=168). In this way we aimed to identify the transcriptional changes associated with HCC malignant transformation and compare these with our findings in murine methyl donor deficiency.

**Fig. 6.**
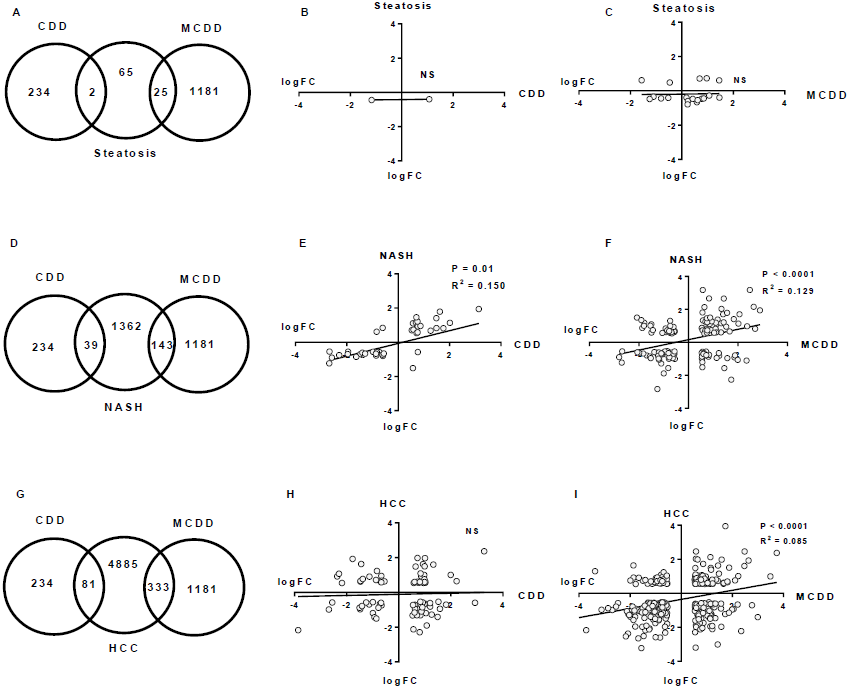
Cross-species comparison of CDD and MCDD transcriptional changes with human stages of NAFLD. Venn diagrams showing common and distinct dysregulated gene orthologues between CDD and MCDD livers and human hepatic steatosis (A), human NASH (D) and human HCC (G). Scatter plots demonstrate Log2 fold change in common dysregulated orthologues between CDD and MCDD mouse livers and corresponding orthologues in human steatosis (B+C), human NASH (E+F) and human HCC (H+I). Trend line, P value and R^2^ value calculated by linear regression analysis.

The CDD and MCDD transcriptomes demonstrated very limited similarity to all stages of human NAFLD. Only 2 (3%) of genes identified as dysregulated in human steatosis were also dysregulated in CDD livers (Fig. 6A and B). 26 (40%) of genes identified as dysregulated in human steatosis were also altered in MCDD mice, however changes in expression of the most commonly dysregulated genes were not in the same direction and there was no significant correlation on linear regression analysis (P = 0.9) (Fig. 6C). There was a greater but still comparatively small overlap in dysregulated gene sets from human NASH studies and CDD and MCDD mice (39 (2.8%) and 143 (10.4%) respectively (Fig. 6D). Those transcripts that were dysregulated in NASH and CDD or MCDD mice did demonstrate a weak but significant correlation in terms of directional change (P 0.01, R^2^ 0.150 for CDD and P < 0.001, R^2^ 0.129 for MCDD) (Fig. 6E and F). Upregulated transcripts common to both CDD mice and NASH were almost exclusively involved in inflammatory *(Lgals3*, *Cd52, Clec7a*) and malignant processes (Tm4sf4, *S100A11, Gpnmb)*. These genes were also upregulated in MCDD and NASH, in which there were additional changes in fibrosis regulators (*Lum*, *Osbpl3*, *Col6a3*, *Tgfbi*, *Tmsb10*, *Tpm1*) and oncogenes (*Golm1* and *Emp1*). Down-regulated genes common to CDD, MCDD and NASH were overrepresented in the GO terms “GO:0006629 Lipid metabolic process” and included the master lipid regulator *Mlxipl* and the lipid synthesis enzymes *Acat2, Agpat2, Lss, Acacb* and *Mvd.*

When comparing the transcriptome of each dietary intervention to HCC, there was again only minimal overlap in perturbed transcripts (81 (1.6%) in CDD and 333 (7.6%) in MCDD) (Fig. 6G). There was no correlation in terms of directional change between CDD and HCC (Fig. 6H). Overlapping transcripts between MCDD mice and HCC did show a weak and highly significant agreement in direction of transcriptional change (R^2^ 0.085, P < 0.0001, Fig. 6I). Genes which were dysregulated in both MCDD and CDD datasets and in HCC were overrepresented in GO terms ‘GO:0000278 mitotic cell cycle’, ‘GO:0007599 Haemostasis’ and ‘GO:0070373 negative regulation of ERK1 and ERK2 cascade’ and include putative HCC oncogenes *Cdc20*, *Osgin1* and *Cdk1*.

## Discussion

It is widely assumed that the steatosis induced by CDD and MCDD results from impaired export of VLDLs, which are required for triglyceride clearance from hepatocytes, perhaps because deficiency of choline and methionine results in an inability to synthesise the major lipid bilayer component phosphatidylcholine (PC) required for VLDL synthesis ^30,34^. Our study supports the concept that both decreased PC synthesis and impaired VLDL secretion may play a role in the hepatic pathology in these models and suggest a potential role for the carboxylesterase (Ces) enzymes in mediating the reduction in VLDL secretion.

The importance of reduced hepatic lipid clearance in MCDD is supported by studies demonstrating i) reduced clearance of radiolabelled hepatic fatty acids, ii) a decrease in serum VLDL concentrations and iii) reduced serum triglyceride accumulation in the context of the peripheral lipase inhibitor typoxalol ^17,26^. Additionally, increased hepatic sequestration of radiolabelled fatty acids and increased incorporation of ^14^C into hepatic triglycerides suggest that increased lipid uptake and/or increased *de novo* lipogenesis may also occur with MCDD ^17,26,34^. These effects have not been reported in rodents exposed to CDD alone ^17,35^; indeed *ex vivo* studies using primary hepatocytes isolated from rats maintained on CDD has shown that the presence of methionine is sufficient to maintain normal levels of PC synthesis and VLDL export into culture media ^34^ and similar experiments in mouse primary hepatocytes demonstrated only a minor reduction in triglyceride export and no change in apolipoprotein secretion in choline deficient media ^9^. These findings may be due to the presence of an accessory pathway for PC synthesis which is only present in liver, where in the absence of choline, PC can be directly synthesised from phosphatidylethanolamine (PE) by the enzyme phosphatidylethanolamine N-methyltransferase (PEMT) using methionine as a methyl donor. Indeed ~30% of PC is synthesised in this way in rodent liver ^36^.

In our study, detailed analysis of the expression of genes in *de novo* lipogenesis pathways in CDD and MCDD strongly suggest an appropriate compensatory response to the high hepatic triglyceride content, with a clear suppression of key mediators of fatty acid synthesis and elongation and cholesterol synthesis (Figure 7). This supports the concept that impaired lipid clearance rather than impaired *de novo* lipogenesis is responsible for the hepatic fat accumulation that occurs with both diets. Consistent with this, the expression of the Ces enzymes (Ces1d, Ces1f, Ces3b) was markedly suppressed in MCDD and the expression of the liver predominant Ces isoform, Ces3b, was also suppressed in CDD. These enzymes are important regulators of VLDL lipid packaging and assembly in the hepatic endoplasmic reticulum (ER), and as such a reduction in expression would be expected to result in reduced hepatic lipid clearance ^20^. Mice lacking liver specific Ces3 (also known as triacylglycerol hydrolase) have a reduction in circulating VLDL triglycerides and cholesterol levels on a standard chow diet with altered hepatic lipid droplet morphology ^20,37^. Furthermore, Ces1 overexpression in mice reduces hepatic triglyceride content and plasma glucose levels whereas liver specific knock-down results in increased hepatic triglyceride ^19^. Whilst it is unclear why the expression of these genes is suppressed in the presence of an increased hepatic lipid load (notably in MCDD), we suggest that these models may present an opportunity for investigating the mechanism of action of these important hepatic lipid clearance enzymes and the screening of therapeutics that exploit these molecular targets. The expression of Patatin-like phospholipase domain containing 3 (Pnpla3) was also suppressed in both CDD and MCDD models. The human PNPLA3^I148M^ variant is strongly associated with human NAFLD ^38^; humans homozygous for the PNPLA3^I148M^ allele are reported to have ~73% more hepatic triglyceride when compared with matched heterozygote controls, and both *in vivo* and *in vitro* studies suggest that this is due to impaired triglyceride hydrolysis and VLDL export ^39–42^. Thus, Pnpla3 suppression may also contribute to triglyceride accumulation in methyl donor deficiency.

**Fig. 7.**
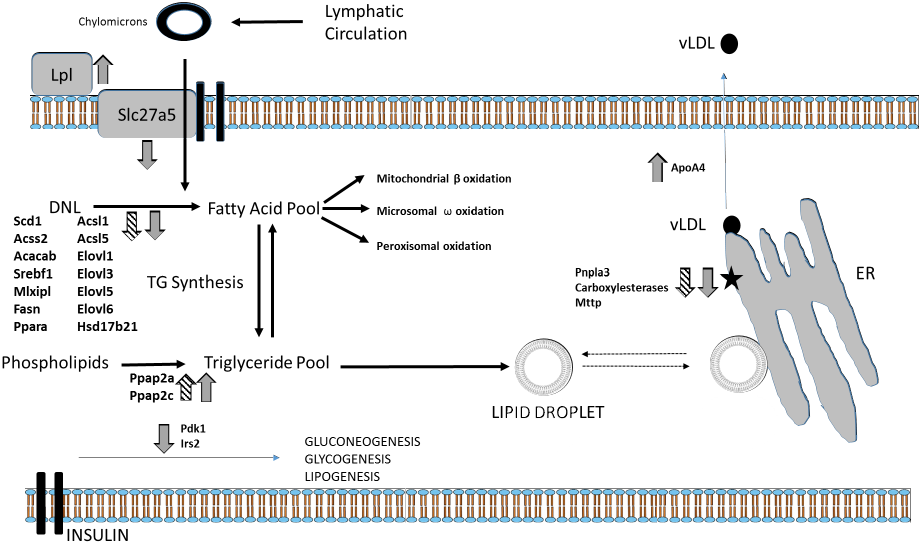
Transcriptional dysregulation mapped to pathways of lipid transport and metabolism in hepatocytes. Arrows demonstrate direction of transcriptional change isn CDD (hatched) and MCDD (grey) diets. *Pnpla3* and *Ces* suppression suggest impaired packaging of lipid into VLDL particles on the surface of the ER (black star), which may represent a novel site of lipid accumulation.

Dissection of the interacting pathways involved in choline, methionine and PC metabolism in these models provides further insights into the mechanisms by which the murine liver responds to dietary choline and methionine deficiency. Whilst choline has a major role as a substrate for PC synthesis ^36^, the essential amino acid methionine is also necessary for the methylation of a large variety of substrates including DNA, proteins and lipids and for the synthesis of polyamines, and it is also crucial for normal hepatocyte function ^43^. Both substrates are important for the maintenance of hepatic SAMe levels, which are normally tightly regulated to maintain normal hepatic function ^15^, and the direction of transcriptional changes with choline and methoinine deficiency strongly suggest a drive to maintain hepatic SAMe concentrations. The down-regulation of Chkb and Pcyta which are involved in the synthesis of PC from choline, coupled with the upregulation of methionine adenosyl-transferase (Mat2a), which synthesises SAMe from methionine, suggest a forward drive to maintain SAMe levels. The concurrent down-regulation of Gnmt and Ahcy (which metabolise SAMe and SAH respectively) may act as a further cellular buffer to maintain SAMe concentrations ^44^. Nevertheless, despite these changes, we found reduced levels of hepatic SAMe in MCDD mice in agreement with other studies ^45^, suggesting an inability to maintain SAMe leveles with severe deficiency of both substrates. Thus, the cumulative effect of the observed transcriptional changes in MCDD mice is directed at maintaining SAMe concentrations at the expense of PC synthesis, with the potential to result in decreased VLDL synthesis. Further evidence in support of the importance of SAMe deficiency in the pathogenesis of liver disease in MCDD mice is supported by the fact that the deleterious effects of MCDD diets can be rescued by the administration of SAMe ^46^.

Whilst there are some clear biological similarities between the hepatic pathology induced by methyl donor deficiency in rodents and human NAFLD/NASH ^47–49,35,50^, there are also a number of major differences. In humans, NAFLD is closely associated with obesity and insulin resistance, whereas in rodents, CDD results in profound hepatic steatosis without insulin resistance ^17,51^ and MCDD causes an inflammatory steatohepatitis with fibrogenesis and significant weight loss with an *increase* in peripheral insulin sensitivity ^30,52^. Our transcriptomic analysis also suggests that CDD and MCDD produce a hepatic phenotype which is markedly dissimilar to human NAFLD in terms of lipid handling. Whereas human NAFLD is associated with an upregulation of genes important in *de novo* lipogenesis (FASN, MLPXL, ACACA, SREB-1c) ^53,54^, this is either not seen, or indeed the reverse is observed in mice maintained on CDD/MCDD diets. Furthermore, although some findings in human NASH support the concept that NAFLD may result at least in part in from an inability to synthesise PC ^55,56^, none of the genes dysregulated in the one carbon metabolism pathways of interest in CDD/MCDD were also altered in the human simple steatosis or NASH data sets. Although SAMe depletion is a feature of human NAFLD and correlates with severity of disease in NAFLD biopsies, and oral SAMe preparations are currently under review as a treatment for chronic liver disease ^14,16^, mediators of SAMe metabolism were not altered the human statosis or NASH datasets. Some enzymes important in SAMe metabolism were dysregulated in the human HCC data (upregulated AHCY (LogFC 0.58); down-regulated BHMT (logFC-1.35), GNMT (LogFC −1.05) and MAT1a (logFC −0.97)), however these did not reflect the changes seen in the mouse model apart from a similar change in the expression of GNMT.

In conclusion, our data suggest a novel alternative mechanism for methyl donor deficient liver injury involving impaired VLDL particle assembly due to suppression of key triglyceride hydrolysis proteins. Although these CDD and MCDD models are widely used for the study of NAFLD, their translational impact in studies of NAFLD/NASH is likely to be limited by fundamental differences in the global transcriptional profiles between these models and human disease states. Our data do suggest that MCDD may be a useful model for studying the development of HCC secondary to the premalignant inflammatory steatohepatitis NASH. We suggest that there remains an urgent need for novel, more representative models of the full spectrum of NAFLD pathology.

## Acknowledgements/grant support

Our thanks go the the Edinburgh Clinical Research Facility Genetics Core, Western General Hospital, Edinburgh, UK for technical expertise with microarray processing.

---

**Supplementary Figure 1. Gene Ontology analysis of CDD and MCDD mouse liver.** Overrepresented pathways of up (black) and down (hatched) dysregulated genes in CDD (A+B) and MCDD (C+D) mouse liver microarray analysis.

**Supplementary Table 1** Dietary constituents

**Supplementary Table 2** Primer sequences

**Supplementary Table 3** Top 100 differentially expressed genes in CDD mice versus control animals.

**Supplementary Table 4** Top 100 differentially expressed genes in MCDD mice versus control animals.

**Supplementary Table 5** Details of subject numbers and arrays used from human studies

## References

1. Loomba, R. & Sanyal, A. J. The global NAFLD epidemic. Nat Rev Gastroenterol Hepatol 10, 686–690 (2013).

2. Leung, C. et al. Characteristics of hepatocellular carcinoma in cirrhotic and non-cirrhotic non-alcoholic fatty liver disease. World J. Gastroenterol. 21, 1189–96 (2015).

3. Smits, M. M., Ioannou, G. N., Boyko, E. J. & Utzschneider, K. M. Non-alcoholic fatty liver disease as an independent manifestation of the metabolic syndrome: results of a US national survey in three ethnic groups. J Gastroenterol Hepatol 28, 664–670 (2013).

4. Perry, R. J., Samuel, V. T., Petersen, K. F. & Shulman, G. I. The role of hepatic lipids in hepatic insulin resistance and type 2 diabetes. Nature 510, 84–91 (2014).

5. Blachier, M., Leleu, H., Peck-Radosavljevic, M., Valla, D. C. & Roudot-Thoraval, F. The burden of liver disease in Europe: a review of available epidemiological data. J Hepatol 58, 593–608 (2013).

6. Ito, M. et al. Longitudinal analysis of murine steatohepatitis model induced by chronic exposure to high-fat diet. Hepatol. Res. 37, 50–57 (2007).

7. Gallou-Kabani, C. et al. C57BL/6J and A/J mice fed a high-fat diet delineate components of metabolic syndrome. Obesity (Silver Spring). 15, 1996–2005 (2007).

8. Duval, C. et al. Adipose tissue dysfunction signals progression of hepatic steatosis towards nonalcoholic steatohepatitis in C57Bl/6 mice. Diabetes 59, 3181–3191 (2010).

9. Kulinski, A., Vance, D. E. & Vance, J. E. A choline-deficient diet in mice inhibits neither the CDP-choline pathway for phosphatidylcholine synthesis in hepatocytes nor apolipoprotein B secretion. J. Biol. Chem. 279, 23916–24 (2004).

10. Leclercq, I. A., Lebrun, V. A., Starkel, P. & Horsmans, Y. J. Intrahepatic insulin resistance in a murine model of steatohepatitis: effect of PPARgamma agonist pioglitazone. Lab Invest 87, 56–65 (2007).

11. Lyman, R. L., Giotas, C., Medwadowski, B. & Miljanich, P. Effect of low methionine, choline deficient diets upon major unsaturated phosphatidyl choline fractions of rat liver and plasma. Lipids 10, 157–167 (1975).

12. De L’Hortet, a. C. et al. GH administration rescues fatty liver regeneration impairment by restoring GH/EGFR pathway deficiency. Endocrinology 155, 2545–2554 (2014).

13. Okubo, H. et al. Involvement of resistin-like molecule beta in the development of methionine-choline deficient diet-induced non-alcoholic steatohepatitis in mice. Sci. Rep. 6, 20157 (2016).

14. Martinez-Chantar, M. L. et al. Importance of a deficiency in S-adenosyl-L-methionine synthesis in the pathogenesis of liver injury. Am J Clin Nutr 76, 1177s–82s (2002).

15. Mato, J. M. & Lu, S. C. Role of S-adenosyl-L-methionine in liver health and injury. Hepatology 45, 1306–1312 (2007).

16. Anstee, Q. M. & Day, C. P. S-adenosylmethionine (SAMe) therapy in liver disease: A review of current evidence and clinical utility. J. Hepatol. 57, 1097–1109 (2012).

17. Macfarlane, D. P. et al. Metabolic pathways promoting intrahepatic fatty acid accumulation in methionine and choline deficiency: implications for the pathogenesis of steatohepatitis. Am. J. Physiol. Endocrinol. Metab. 300, E402–409 (2011).

18. Swales, J. G. et al. Mass Spectrometry Imaging of Cassette-Dosed Drugs for Higher Throughput Pharmacokinetic and Biodistribution Analysis. Anal Chem (2014). DOI:10.1021/ac502217r

19. Xu, J. et al. Hepatic carboxylesterase 1 is essential for both normal and farnesoid X receptor-controlled lipid homeostasis. Hepatology 59, 1761–1771 (2014).

20. Lian, J. et al. Liver specific inactivation of carboxylesterase 3/triacylglycerol hydrolase decreases blood lipids without causing severe steatosis in mice. Hepatology 56, 2154–2162 (2012).

21. Quiroga, A. D. et al. Deficiency of carboxylesterase 1/esterase-x results in obesity, hepatic steatosis, and hyperlipidemia. Hepatology 56, 2188–2198 (2012).

22. Ahrens, M. et al. DNA methylation analysis in nonalcoholic fatty liver disease suggests distinct disease-specific and remodeling signatures after bariatric surgery. Cell Metab 18, 296–302 (2013).

23. Lake, A. D. et al. Analysis of global and absorption, distribution, metabolism, and elimination gene expression in the progressive stages of human nonalcoholic fatty liver disease. Drug Metab. Dispos. 39, 1954–60 (2011).

24. Villanueva, A. et al. DNA methylation-based prognosis and epidrivers in hepatocellular carcinoma. Hepatology 61, 1945–1956 (2015).

25. Raubenheimer, P. J., Nyirenda, M. J. & Walker, B. R. A choline-deficient diet exacerbates fatty liver but attenuates insulin resistance and glucose intolerance in mice fed a high-fat diet. Diabetes 55, 2015–2020 (2006).

26. Rinella, M. E. et al. Mechanisms of hepatic steatosis in mice fed a lipogenic methionine choline-deficient diet. J. Lipid Res. 49, 1068–1076 (2008).

27. Ip, E. et al. Central role of PPARalpha-dependent hepatic lipid turnover in dietary steatohepatitis in mice. Hepatology 38, 123–132 (2003).

28. Johnson, E. F., Hsu, M.-H., Savas, U. & Griffin, K. J. Regulation of P450 4A expression by peroxisome proliferator activated receptors. Toxicology 181–182, 203–206 (2002).

29. Leclercq, I. A. et al. CYP2E1 and CYP4A as microsomal catalysts of lipid peroxides in murine nonalcoholic steatohepatitis. J. Clin. Invest. 105, 1067–1075 (2000).

30. Rinella, M. E. & Green, R. M. The methionine-choline deficient dietary model of steatohepatitis does not exhibit insulin resistance. J Hepatol 40, 47–51 (2004).

31. Rubio, A. et al. Identification of a gene-pathway associated with non-alcoholic steatohepatitis. J. Hepatol. 46, 708–718 (2007).

32. Younossi, Z. M. et al. A genomic and proteomic study of the spectrum of nonalcoholic fatty liver disease. Hepatology 42, 665–674 (2005).

33. Starmann, J. et al. Gene expression profiling unravels cancer-related hepatic molecular signatures in steatohepatitis but not in steatosis. PLoS One 7, e46584 (2012).

34. Yao, Z. M. & Vance, D. E. The active synthesis of phosphatidylcholine is required for very low density lipoprotein secretion from rat hepatocytes. J Biol Chem 263, 2998–3004 (1988).

35. Veteläinen, R., van Vliet, A. & van Gulik, T. M. Essential pathogenic and metabolic differences in steatosis induced by choline or methione-choline deficient diets in a rat model. J Gastroenterol Hepatol 22, 1526–1533 (2007).

36. >Li, Z. & Vance, D. E. Phosphatidylcholine and choline homeostasis. J. Lipid Res. 49, 1187–1194 (2008).

37. Wang, H. et al. Altered lipid droplet dynamics in hepatocytes lacking triacylglycerol hydrolase expression. Mol. Biol. Cell 21, 1991–2000 (2010).

38. He, S. et al. A sequence variation (I148M) in PNPLA3 associated with nonalcoholic fatty liver disease disrupts triglyceride hydrolysis. J. Biol. Chem. 285, 6706–15 (2010).

39. Pirazzi, C. et al. Patatin-like phospholipase domain-containing 3 (PNPLA3) I148M (rs738409) affects hepatic VLDL secretion in humans and in vitro. J. Hepatol. 57, 1276–82 (2012).

40. Huang, Y., Cohen, J. C. & Hobbs, H. H. Expression and characterization of a PNPLA3 protein isoform (I148M) associated with nonalcoholic fatty liver disease. J. Biol. Chem. 286, 37085–93 (2011).

41. Chamoun, Z., Vacca, F., Parton, R. G. & Gruenberg, J. PNPLA3/adiponutrin functions in lipid droplet formation. Biol. Cell 105, 219–33 (2013).

42. Sookoian, S. et al. Epigenetic regulation of insulin resistance in nonalcoholic fatty liver disease: impact of liver methylation of the peroxisome proliferator-activated receptor gamma coactivator 1alpha promoter. Hepatology 52, 1992–2000 (2010).

43. Mato, J. M., Martinez-Chantar, M. L. & Lu, S. C. S-adenosylmethionine metabolism and liver disease. Ann. Hepatol. 12, 183–189 (2013).

44. Luka, Z., Mudd, S. H. & Wagner, C. Glycine N-methyltransferase and regulation of S-adenosylmethionine levels. J. Biol. Chem. 284, 22507–22511 (2009).

45. Caballero, F. et al. Specific contribution of methionine and choline in nutritional nonalcoholic steatohepatitis: Impact on mitochondrial S-adenosyl-L-methionine and glutathione. J. Biol. Chem. 285, 18528–18536 (2010).

46. Dahlhoff, C. et al. Methyl-donor supplementation in obese mice prevents the progression of NAFLD, activates AMPK and decreases acyl-carnitine levels. Mol. Metab. 3, 565–80 (2014).

47. Schattenberg, J. M., Wang, Y., Singh, R., Rigoli, R.M. & Czaja, M. J. Hepatocyte CYP2E1 overexpression and steatohepatitis lead to impaired hepatic insulin signaling. J Biol Chem 280, 9887–9894 (2005).

48. Emery, M. G. et al. CYP2E1 activity before and after weight loss in morbidly obese subjects with nonalcoholic fatty liver disease. Hepatology 38, 428–435 (2003).

49. Weltman, M. D., Farrell, G. C., Hall, P., Ingelman-Sundberg, M. & Liddle, C. Hepatic cytochrome P450 2E1 is increased in patients with nonalcoholic steatohepatitis. Hepatology 27, 128–133 (1998).

50. Fisher, C. D. et al. Hepatic cytochrome P450 enzyme alterations in humans with progressive stages of nonalcoholic fatty liver disease. Drug Metab Dispos 37, 2087–2094 (2009).

51. Raubenheimer, P. J., Nyirenda, M.J. & Walker, B. R. A choline-deficient diet exacerbates fatty liver but attenuates insulin resistance and glucose intolerance in mice fed a high-fat diet. Diabetes 55, 2015–2020 (2006).

52. Rizki, G. et al. Mice fed a lipogenic methionine-choline-deficient diet develop hypermetabolism coincident with hepatic suppression of SCD-1. J Lipid Res 47, 2280–2290 (2006).

53. Auguet, T. et al. Liver lipocalin 2 expression in severely obese women with non alcoholic fatty liver disease. Exp Clin Endocrinol Diabetes 121, 119–124 (2013).

54. Higuchi, N. et al. Liver X receptor in cooperation with SREBP-1c is a major lipid synthesis regulator in nonalcoholic fatty liver disease. Hepatol. Res. 38, 1122–9 (2008).

55. Li, Z. et al. The ratio of phosphatidylcholine to phosphatidylethanolamine influences membrane integrity and steatohepatitis. Cell Metab 3, 321–331 (2006).

56. Arendt, B. M. et al. Nonalcoholic fatty liver disease is associated with lower hepatic and erythrocyte ratios of phosphatidylcholine to phosphatidylethanolamine. Appl Physiol Nutr Metab 38, 334–340 (2013).

